# Validating a high-throughput tracking system: ATLAS as a regional-scale alternative to GPS

**DOI:** 10.1101/2021.02.09.430514

**Authors:** Christine E. Beardsworth, Evy Gobbens, Frank van Maarseveen, Bas Denissen, Anne Dekinga, Ran Nathan, Sivan Toledo, Allert I. Bijleveld

## Abstract

1. Fine-scale tracking of animal movement is important to understand the proximate mechanisms of animal behaviour. While GPS tracking is an excellent tool for measuring animal movement, trade-offs between tag weight, cost and lifespan limit its application to relatively large species, a small number of individuals or short tracking durations, respectively. The reverse-GPS system – ATLAS – uses lighter, cheaper tags compared to GPS tags, that can also last long periods of time at high sampling frequencies. Six systems are now operational worldwide and have successfully tracked over 50 species in various landscape types. This growing use of ATLAS to track animal movement motivates further refinement of best-practice application and an assessment of its accuracy.
2. Here, we test the accuracy and precision of the largest ATLAS system, located in the Dutch Wadden Sea using concurrent GPS measurements as a reference. This large-scale ATLAS system consists of 26 receivers and covers 1326 km^2^ of intertidal region, with almost no physical obstacles for radio signals, providing a useful baseline for other systems. To measure accuracy, we calculated the distance between ATLAS and GPS location estimates for a route (mobile test) and 16 fixed locations (stationary test) on the Griend mudflat.
3. ATLAS-derived location estimates differed on average 4.2 m from GPS-estimated stationary test sites and 5.7 m from GPS tracks taken whilst moving between them. Signals that were collected by more receiver stations were more accurate, although even 3-receiver localisations were comparable with GPS localisations (∼10 m difference). Higher receiver stations detected the tag at longer distances.
4. Future ATLAS users should consider the height of receivers, their spatial arrangement, density and the movement mode of the study species (e.g., ground-dwelling or flying). In conclusion, ATLAS provides an accurate, regional-scale alternative to global GPS-based tracking with which hundreds of relatively small-bodied species can be tracked simultaneously for long periods of time. Our study shows that ATLAS is a valid alternative, providing comparable location estimates to GPS.

## Introduction

Tracking animal movements is important to understand the mechanisms underlying animal behaviour, with broad applications for studying key ecological and evolutionary processes (Nathan et al., 2008). Since the advent of tracking technology, insight into the often cryptic movements of animals has helped us to understand behaviours that were previously almost impossible, from the migration patterns of whales (Abrahms et al., 2019) and birds (Gill et al., 2009) to identifying differences between individuals in foraging strategies (Bijleveld et al., 2016; Harris et al., 2020) or confirming the cognitive mechanisms that underlie behaviour in free-ranging individuals (Beardsworth et al., *In press*; Toledo et al., 2020). In recent decades, the rapid development of global navigation satellite systems, including GPS, has led to an explosion in the number of movement ecology studies (Holyoak et al., 2008; Joo et al., 2020). However, while GPS has become one of the primary tools of choice for monitoring movement (Cagnacci et al., 2010; Hebblewhite & Haydon, 2010; Kays et al., 2015), trade-offs between sampling frequency, battery size, tag weight, cost and lifespan can limit its application. Fortunately, alternative systems for tracking animal movement are continuously being developed in an attempt to address these challenges (Williams et al., 2020). These solutions form a diverse toolset for biologging, ranging from autonomous underwater videography (Hawkes et al., 2020) to using biomarkers from tail hair to detect movement through landscapes (Kabalika et al., 2020). Despite these advances, few technologies can attain the fine-scale tracking of movement paths which are a necessity for identifying decision points of individuals (Collet et al., 2017; Harel et al., 2016) or groups (Strandburg-Peshkin et al., 2015), or correlating movements with precise environmental covariates (Eikelboom et al., 2020). One potential alternative for regional-scale studies is ATLAS (Advanced Tracking and Localisation of Animals in real-life Systems; Toledo et al., 2020), a high-throughput system that uses low-cost and lightweight radio-transmitters to track animals at a regional scale.

Radio telemetry has a rich history for use within animal tracking (Amlaner & Macdonald, 1980; Benson, 2010) and traditionally involved using hand-held receivers to search a landscape for signals from animal-mounted, high frequency radio-transmitters to estimate tag location through triangulation. Unlike GPS, radio tags act as transmitters rather than receivers, alleviating energy-demanding position computations and remote data communications. Miniature radio-transmitters have low power requirements (thus requiring smaller batteries), increasing their potential for use with smaller species while keeping cost minimal. However, conventional radio telemetry is labour intensive and it is not feasible to follow more than a few individuals and locations are estimated irregularly. Attempts to automate wildlife tracking based on standard radio tags were made as early as the 1960’s (Cochran et al., 1965) and more recently using a bounding array of receivers distributed within a specific region (Kays et al., 2011). Locations can be estimated from the data collected by these receivers and has the potential to be accessed almost instantaneously. However, while previous automated radio-telemetry systems have been unable to match the accuracy of GPS, ATLAS uses the same location estimation technique as GPS, time of arrival (TOA). Rather than approximating the angle of arrival of the signal (where a 1-degree error becomes an error of 17 m from 1km away), an error in time of arrival of 10ns remains an error of 10ns at all distances. One difficulty of TOA systems is the need for absolutely accurate clocks (MacCurdy et al., 2009), but through implementing the use of beacon tags in known locations to synchronise clocks, ATLAS relies more heavily on stability of clocks than accuracy (Weller-Weiser et al., 2016). Through these beacons, ATLAS can synchronise clocks, characterise the accuracy of localisations and monitor the performance of the system. ATLAS therefore provides high-throughput monitoring for relatively cheap (∼€50 per tag), lightweight (<1g + battery weight), and long-lasting (∼8 months for 4.4g tag at 1/6 Hz) tags.

However, ATLAS installation requires time, resources and expertise, and its spatial coverage is limited to a regional scale where the line-of-sight from three or more receiver stations overlap. Considering these pros and cons, six ATLAS systems have recently been established in four different countries (Israel, the UK, the Netherlands and Germany), collecting rich datasets on hundreds of individuals of over 50 different species, and addressing key research questions in movement ecology (Beardsworth et al., In press; Corl et al., 2020; Toledo et al., 2020). Yet detailed, strategic testing of the accuracy of ATLAS and the efficacy of receiver arrays has been limited. Here, we characterise the accuracy and precision of the largest ATLAS system to date. The Wadden Sea ATLAS system (WATLAS) has 26 receiver stations covering a total area of 1326 km^2^ which are most concentrated on the Griend mudflat, an important stop-over site for many shorebird species (Piersma et al., 1993). With almost no interruptions to the flat landscape of the intertidal zone, this system is particularly useful to investigate the efficacy of ATLAS. We assess tag reception capabilities and the influence of receiver arrangement and concentration. We compare ATLAS localisations to GPS-derived positions at 16 test sites covering an area of 34 km^2^ around the Griend mudflat. While stationary tests give a good overview of the accuracy and precision of a system, animals themselves are tracked in both stationary and mobile states. It is therefore useful to compare GPS and ATLAS while moving across a landscape, where slight variations in the array of receivers that detect the tag may occur. We calculated the difference between ATLAS and GPS en-route to each of the test sites giving us estimates for accuracy in both stationary and moving states. We emphasise that the accuracy of neither GPS nor ATLAS localisations were tested against independent (“known”) finer-scale locations, but were compared to each other. Hence, these comparisons should not be inferred as if ATLAS deviations from GPS measurements represent deviations from the true location (or vice versa), but whether these two methods are comparably accurate and precise. Finally, since movement studies frequently filter and/or smooth movement data to reduce error in location estimates (Lewis et al., 2007), and more importantly, since GPS data are provided only after intense filtering and smoothing (Kaplan & Hegarty, 2005), we assessed the accuracy of the ATLAS system both with and without applying a simple filter-smoothing process.

## Methods

### WATLAS - The Wadden Sea ATLAS System

The WATLAS system consist of 26 ATLAS receiver stations (Fig. 1). Components include a small computer (Intel NUC mini PC running dedicated ATLAS software, *v2020-04-19-stable*), a USRP N200 radio with a WBX40 daughterboard and a GPS disciplined oscillator (GPSDO), which were housed in custom-made watertight containers. The system time/GPSDO of a receiver station is synchronized using the atomic clocks from a GPS unit (which is connected to the USRP N200 radio) and calibrated using seven beacons tags, placed at known locations throughout the study area. Receiver stations were placed around the western Dutch Wadden sea to gain high coverage for tracking migrant shorebirds while they stopover in the area (Fig. 1). Eleven receivers were built on temporary scaffolds on the mudflats and were powered with four 100 W solar panels and a 100 W wind turbine (Ampair) connected to three (105 Ah) batteries. The remaining 15 receivers were installed in places such as on buildings and other stable structures where power was available. Each receiver had a UHF antenna connected to the radio through a custom-built front-end unit (CircuitHub) and a custom built Low-Noise Amplifier. Radio-frequency samples from tags are processed by the receiver’s computer to estimate the arrival times of the signal. All receiver stations were connected to internet using a 3G cellular model (USB dongles, Huawei E3372) to send detection reports to a central server situated at the NIOZ. In real-time, the server calculated location estimates and stored these in a MySQL (v5.7, https://www.mysql.com/) database.

**Figure 1.**
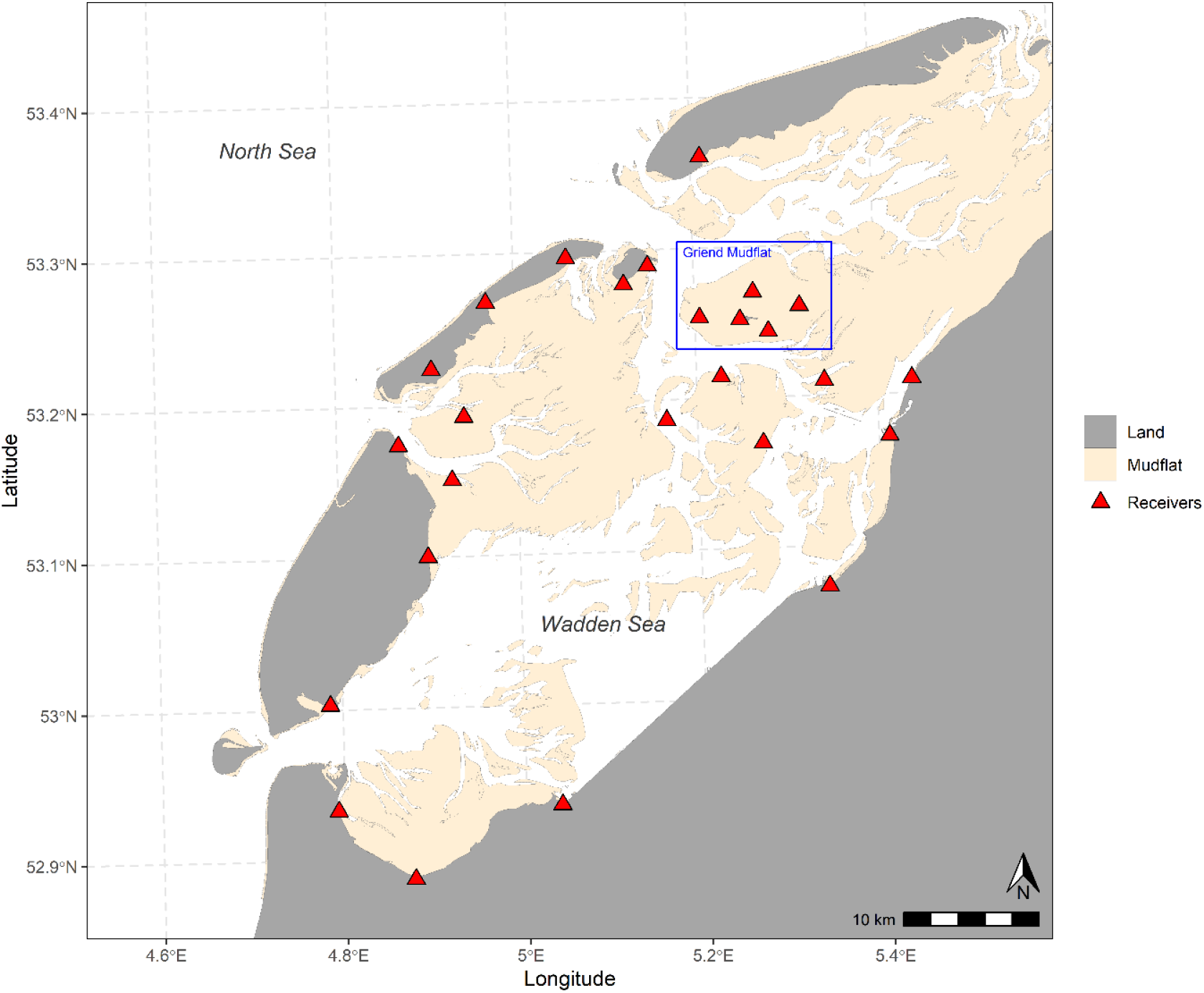
Configuration of receiver stations in 2020 around the Western Wadden sea. The blue rectangle indicates the island of Griend, whose surrounding mudflat provides rich foraging opportunities for many shorebird species and where our receivers are most concentrated. Land is shown in grey and the exposed mudflat at −144 NAP is shown in beige. The locations of receiver stations are shown as red triangles.

### Data collection

To test the accuracy of the WATLAS deployment and ATLAS tracking in general, we focussed our tests on the area around Griend, where we study shorebird movement and have the highest concentration of receivers (Fig. 1). We tested reception and localisation accuracy at 16 sites around the mudflat, 9 of which were 1 km apart with an outer ring of 7 sites that were 2 km between each other and the 1 km sites (Fig. 2). Between 21^st^ - 27^th^ August 2020, we travelled to these sites while carrying a handheld Garmin Dakota 10 GPS (<10 m error 95% typical) – set to record tracks on ‘auto’ which records at a variable rate to create an optimum representation of tracks - and an ATLAS tag. The tag we used for testing emitted a radio signal at 1 Hz. It consisted of a miniature frequency-shift-keying 434MHz integrated radio transceiver and microcontroller (Texas Instruments CC1310) and a monopole ¼ λ gold-plated, multistranded steel wire antenna (Toledo et al., 2014). The tag was encased in plastic to protect it from mechanical stress. We attached the tag to the top of a 1.2m wooden pole so that we could keep the height of the tag consistent throughout the testing.

**Figure 2.**
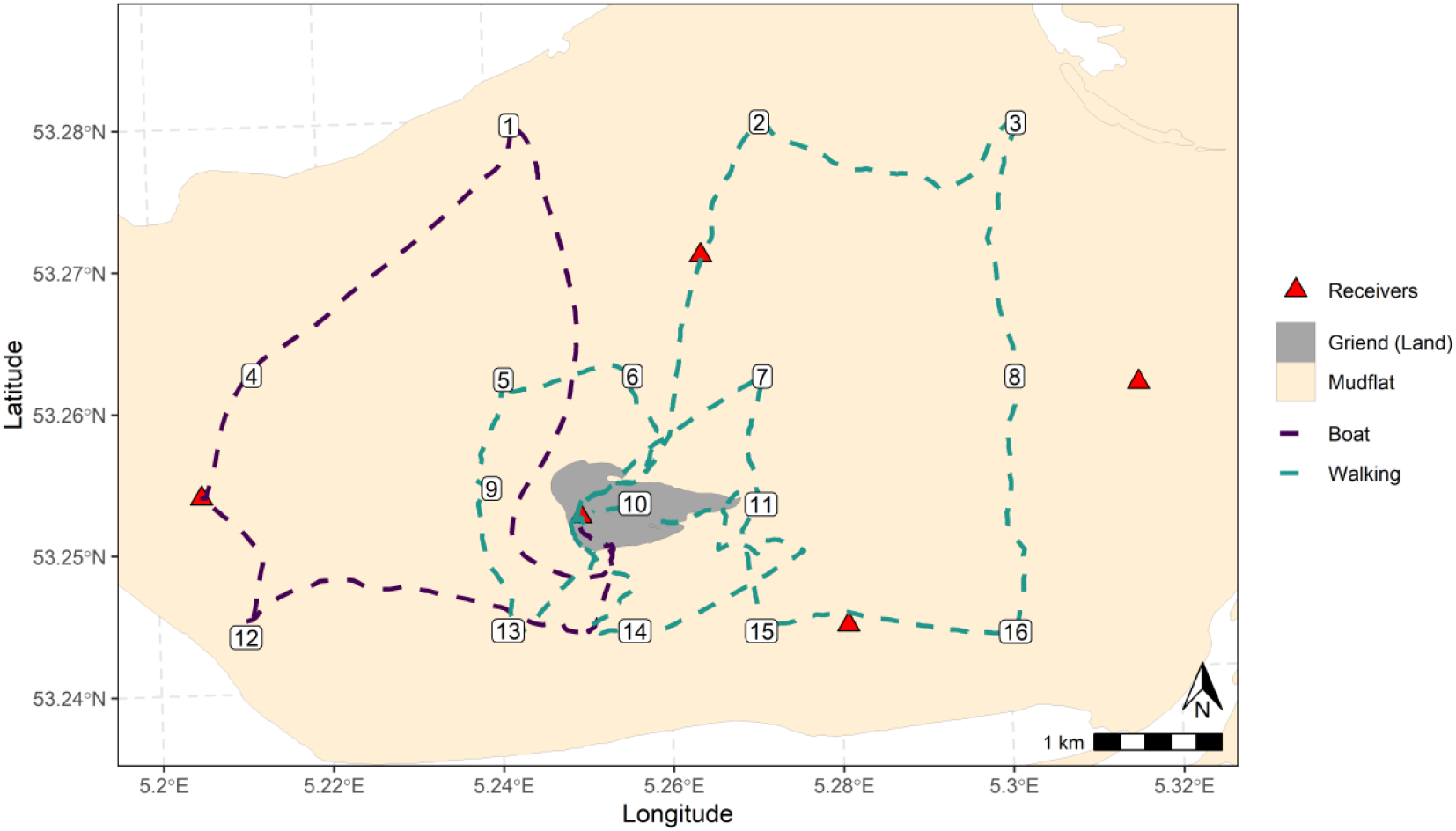
Stationary test sites (numbered) on the Griend mudflat and routes between them which are mapped using GPS data. Due to time and weather constraints, we used a boat to test the western side of the mudflat (purple) but the rest was walked (green).

We walked to 13 of the sites during the day at low tide on the 21^st^, 23^rd^ and 27^th^ August (Fig. 2). Between sites, the pole holding the tag was kept upright meaning that the tag was consistently ∼1.2m from the ground. On arrival at each of these sites, we pushed the pole into the sand so that the tag was 1m above the mudflat and attached the GPS to the pole. We collected ATLAS and GPS data at each site for 5 minutes. Due to weather and time constraints, we were unable to travel to each site on foot. On the 24^th^ August, we sailed a rubber boat to the some of the furthest sites in the western and northern mudflat (Fig. 2) and conducted the stationary test for sites 1, 4 and 12 while at anchor. The pole with the ATLAS tag and GPS tag attached was held upright on the pole in the middle of the boat therefore the tag was ∼1.3m above the water level. Despite being at anchor with a taut rope, there were waves and therefore the boat was not completely stationary during the test, therefore we expect slight overestimates in the location error and larger standard deviations for these positions.

### Filter-Smoothing

Many biologists choose to filter and/or smooth their data before use to reduce errors in positioning estimates (Bjørneraas et al., 2010; Gupte et al., 2020). As GPS data are regularly smoothed without a practical option to retrieve the raw data (Kaplan & Hegarty, 2005), we used a simple filter-smoothing process on the raw data. ATLAS provides some error estimates automatically, namely variance in the Easting and Northing (VARX and VARY). We removed localisations that had high VARX and VARY (>2000) and then smoothed the data by computing a 3-point median smooth across the localisations (Appendix S1).

### Accuracy Analysis and Tag Reception

We split the analysis into two parts, analysing data at test sites (stationary test) and between test sites (mobile test) separately.

#### Stationary Test

For the stationary test, we calculated the mean GPS-derived location estimate at each of the 16 test sites and compared it to the ATLAS-derived location estimate; hence, our measure of accuracy (which we henceforth call ‘error’) refers to the difference in metres between the two estimated locations. For each site, the mean error (m), standard deviation, the median error (m) and the 95^th^ upper and lower percentiles. We investigated the tag reception at each location by calculating the fix rate (number of localisations/300 (max possible localisations in 5-minute period at 1Hz) and the mean and standard deviation of the number of receivers that contributed to each location estimate.

#### Mobile Test

When the tag was mobile, we compared each ATLAS-derived location estimate to the nearest (in time) GPS-derived location estimate. We removed pairings of location estimates where the smallest time difference between an ATLAS location estimate and a GPS location estimate was >2 seconds. We calculated the distance (m) between the remaining paired locations to determine ‘error’. As in the stationary test, we calculated the mean error (m), standard deviation, the median error (m) and the 95^th^ upper and lower percentiles to assess accuracy. We aggregated these summary statistics by the number of receivers that contributed to each location estimate to assess the influence of the number receiver stations on accuracy and investigate the overall coverage of our system. To assess tag reception around the mudflat for specific receivers, we plotted each receiver station and the location estimates that they contributed to on separate maps.

## Results

### Stationary Test

For the stationary test, we calculated a median error [2.5-97.5%] of 4.2 m [0.6-110.1] over all sites, which reduced to a median 3.1 m [0.5-27.5] after filter-smoothing the data (Fig. 3). When analysed individually, we found that the least accurate sites were the three most Northern sites (1-3). These sites were outside of the array of receivers and each had a mean of <4 receiver stations detecting them (Fig. 3, Table 1). Fifteen out of sixteen sites had a fix rate of >90%. One site (site 2) had a fix rate of 73% and the lowest accuracy (median error = 110.5 m [5.5-1059.4]) and mean number of receivers detecting the tag there (mean error ± s.d. =3.4 ± 0.8). This low accuracy was able to be mitigated by applying the filter-smooth, which increased accuracy to 28.2 [7.6-67.5]. It should be noted however that for this particular site the number of location estimates that remained after smoothing was only 9.7% of the expected number of points (300).

**Figure 3.**
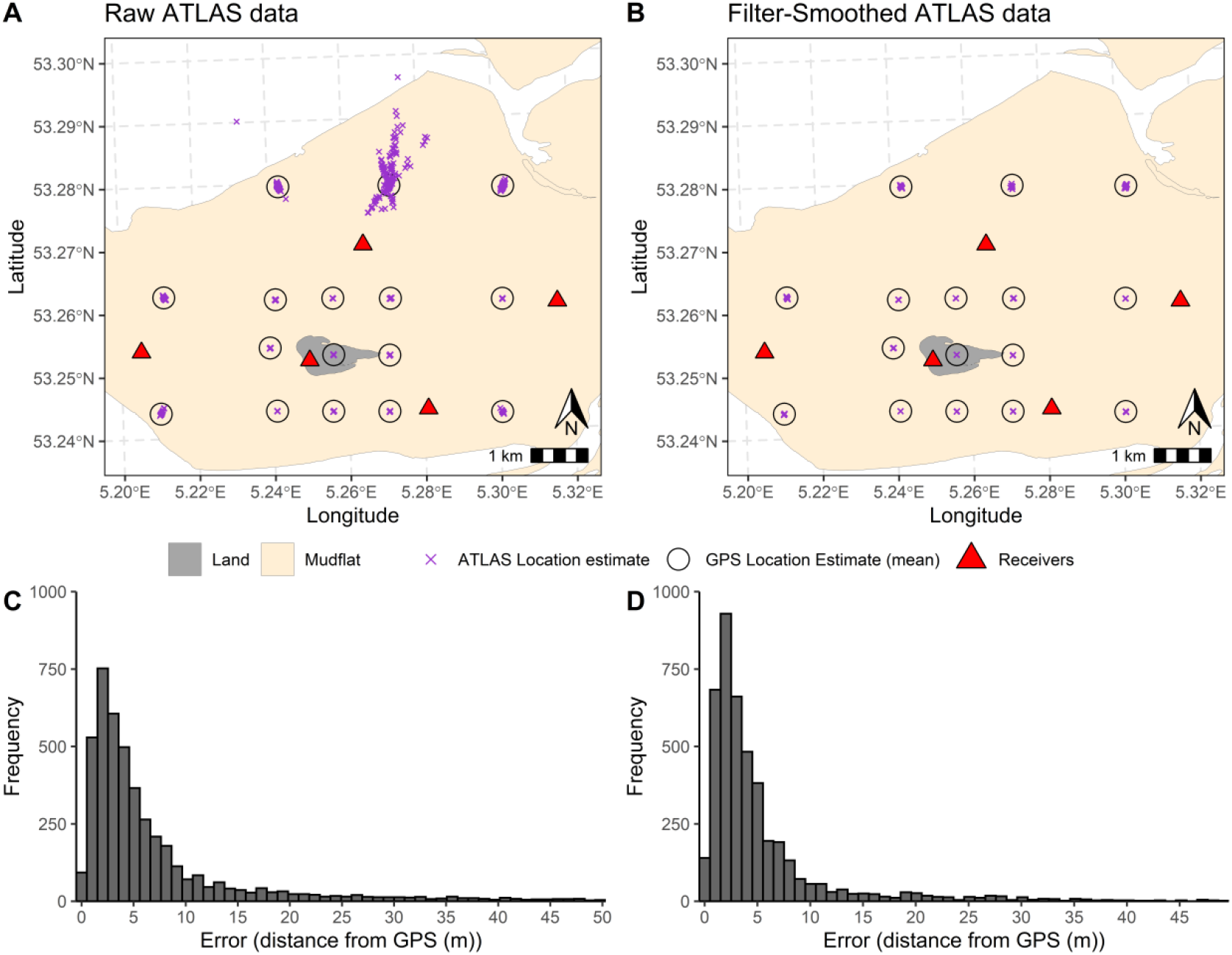
Comparison between ATLAS and GPS derived localisations for raw (A & C) and filter-smoothed (B & D) data. Purple crosses denote positions calculated by ATLAS over a 5-minute period and larger, the centre of black circles indicate the GPS-derived location estimate. The histograms (C & D) show the error in positions at all stationary sites combined, indicating with a peak in accuracy < 5m for both raw and filter-smoothed data.

**Table 1.** Accuracy (error = distance (m) to GPS-derived location estimate) and fix rate (% localisations at 1Hz, max 300) of ATLAS tags (both raw data and filter-smoothed data) that were stationary over five minutes. IDs relate to locations on Fig. 2. ID1, ID4 and ID12 (boxed) were taken on an anchored boat, therefore we may expect higher standard deviations on these results.

| ID | <u>Raw</u> |  |  |  |  | <u>Filter-Smoothed</u> |  |  |  |  |
| --- | --- | --- | --- | --- | --- | --- | --- | --- | --- | --- |
|  | N | Fix Rate (%) | Mean N Receivers (SD) | Mean Error (m) (SD) | Median Error (m) [95%] | N | Fix Rate (%) | Mean N Receivers (SD) | Mean Error (m) (SD) | Median Error (m) [95%] |
| 1 | 276 | 92 | 3.8 (0.8) | 23.8 (83.8) | 13.9 [1.5-59.5] | 265 | 88.3 | 3.8 (0.8) | 11.7 (8.3) | 9.1 [1.2-30.1] |
| 2 | 219 | 73 | 3.4 (0.6) | 248.2 (299.6) | 110.5 [5.5-1059.4] | 29 | 9.7 | 3.6 (0.8) | 28.8 (15.1) | 28.2 [7.6-67.5] |
| 3 | 299 | 99.7 | 3.8 (0.4) | 29.5 (19.9) | 26.3 [4.1-77.6] | 282 | 94 | 3.8 (0.4) | 21 (12.0) | 19.4 [4.3-47.8] |
| 4 | 298 | 99.3 | 3.4 (0.6) | 12.5 (9.9) | 9.4 [1.6-40.6] | 298 | 99.3 | 3.4 (0.6) | 8.2 (6.1) | 7 [1.3-28.5] |
| 5 | 300 | 100 | 4.6 (0.7) | 3.2 (2.4) | 2.9 [0.3-8.5] | 300 | 100 | 4.6 (0.7) | 2.3 (1.5) | 2 [0.3-6.2] |
| 6 | 300 | 100 | 5.3 (0.7) | 3.1 (1.7) | 2.8 [0.7-7.4] | 300 | 100 | 5.3 (0.7) | 2.7 (1.4) | 2.5 [0.8-5.9] |
| 7 | 299 | 99.7 | 3.8 (0.4) | 5.3 (3.8) | 4.1 [0.7-14.3] | 299 | 99.7 | 3.8 (0.4) | 3.4 (2.4) | 2.9 [0.4-8.8] |
| 8 | 300 | 100 | 4 (0.1) | 4.7 (2.3) | 4.4 [1-9.7] | 300 | 100 | 4 (0.1) | 4.2(1.6) | 4.2 [1.6-7.5] |
| 9 | 300 | 100 | 5.7 (0.5) | 3.9 (2.5) | 3.4 [0.7-9.7] | 300 | 100 | 5.7 (0.5) | 3.2 (1.9) | 2.8 [0.7-7.9] |
| 10 | 295 | 98.3 | 3.9 (0.4) | 2.4 (1.7) | 2.2 [0.5-6] | 295 | 98.3 | 3.9 (0.4) | 2.1 (1.1) | 2 [0.5-4.5] |
| 11 | 279 | 93 | 4.8 (1.0) | 3 (2.4) | 2.3 [0.5-9.6] | 279 | 93 | 4.8 (0.4) | 2.1 (1.4) | 1.8 [0.4-5.4] |
| 12 | 281 | 93.7 | 4 (0.9) | 8.1 (11.6) | 5.4 [0.8-24.3] | 274 | 91.3 | 4 (1.0) | 4.6 (3.6) | 3.9 [0.5-16] |
| 13 | 282 | 94 | 5.8 (0.8) | 2.2 (1.3) | 2 [0.3-4.8] | 282 | 94 | 5.8 (0.8) | 1.7 (0.9) | 1.6 [0.2-4] |
| 14 | 300 | 100 | 5.2 (0.4) | 4.8 (3.1) | 4.4 [0.4-12.5] | 300 | 100 | 5.2 (0.4) | 4.3 (2.5) | 4.1 [0.6-9.7] |
| 15 | 300 | 100 | 5.8 (0.4) | 3.6 (2.6) | 3.1 [0.5-9.2] | 300 | 100 | 5.8 (0.4) | 2.7 (1.8) | 2.2 [0.2-7.0] |
| 16 | 299 | 99.7 | 5.2 (0.8) | 7.3 (6.9) | 5.7 [0.9-20.5] | 299 | 99.7 | 5.2 (0.8) | 4.8 (2.8) | 4.6 [0.6-11.3] |

### Mobile Test

ATLAS location estimates had an overall median [2.5-97.5%] of 5.7 m [0.9-63.3] when compared to GPS. In general, localisations where only 3 receiver stations detected the tag were the least accurate and accuracy increased with receiver number (Table 2). Larger errors were more likely to occur on the edges and outside of the receiver array (Fig. 4). The filter-smoothing that we implemented decreased overall error to a median of 4.4 m [0.7-27.4] (Table 2, Fig. 4B). Ten receiver stations received tag signals during the mobile test and ranged from detecting 0.58% of signals to 97.87% (Fig. 5). The furthest distance at which a receiver detected a signal was 14,764 m, but this was the highest of the receivers (44.4 m) and only contributed to 1.17% of localisations. The receivers that detected >90% of signals were within 5 km of their furthest detection.

**Table 2.** Accuracy of tag when moving according to the number of receiver stations that detected the tag. Error is calculated by measuring the distance between the ATLAS localisation and the (temporally) closest GPS localisation (if <2 seconds difference).

| NBS | N | <u>Raw</u> |  | <u>Filter-Smoothed</u> |  |  |
| --- | --- | --- | --- | --- | --- | --- |
|  |  | Mean Error (m) (SD) | Median error [95%] | N (N removed by filters) | Mean Error (m) (SD) | Median error [95%] |
| 3 | 4285 | 24.1 (44.8) | 10.7 [1.3-150.1] | 3850 (-435) | 11.1 (19.0) | 6.3 [1-47.6] |
| 4 | 8712 | 9.9 (15.4) | 5.6[0.9-50.3] | 8464 (-248) | 6.5 (7.5) | 4.3 [0.7-25.9] |
| 5 | 4379 | 5.9 (6.4) | 4.4[0.7-18.9] | 4378 (-1) | 4.9 (4.2) | 3.8 [0.6-15.3] |
| 6 | 1358 | 5.3 (3.9) | 4.2 [0.7-15.8] | 1358 (0) | 4.6 (3.4) | 3.6 [0.6-14.2] |
| 7 | 53 | 4.4 (3.5) | 3.4 [1.1-13.1] | 53 (0) | 3.4 (3.0) | 2.3 [0.8-10.5] |

**Figure 4.**
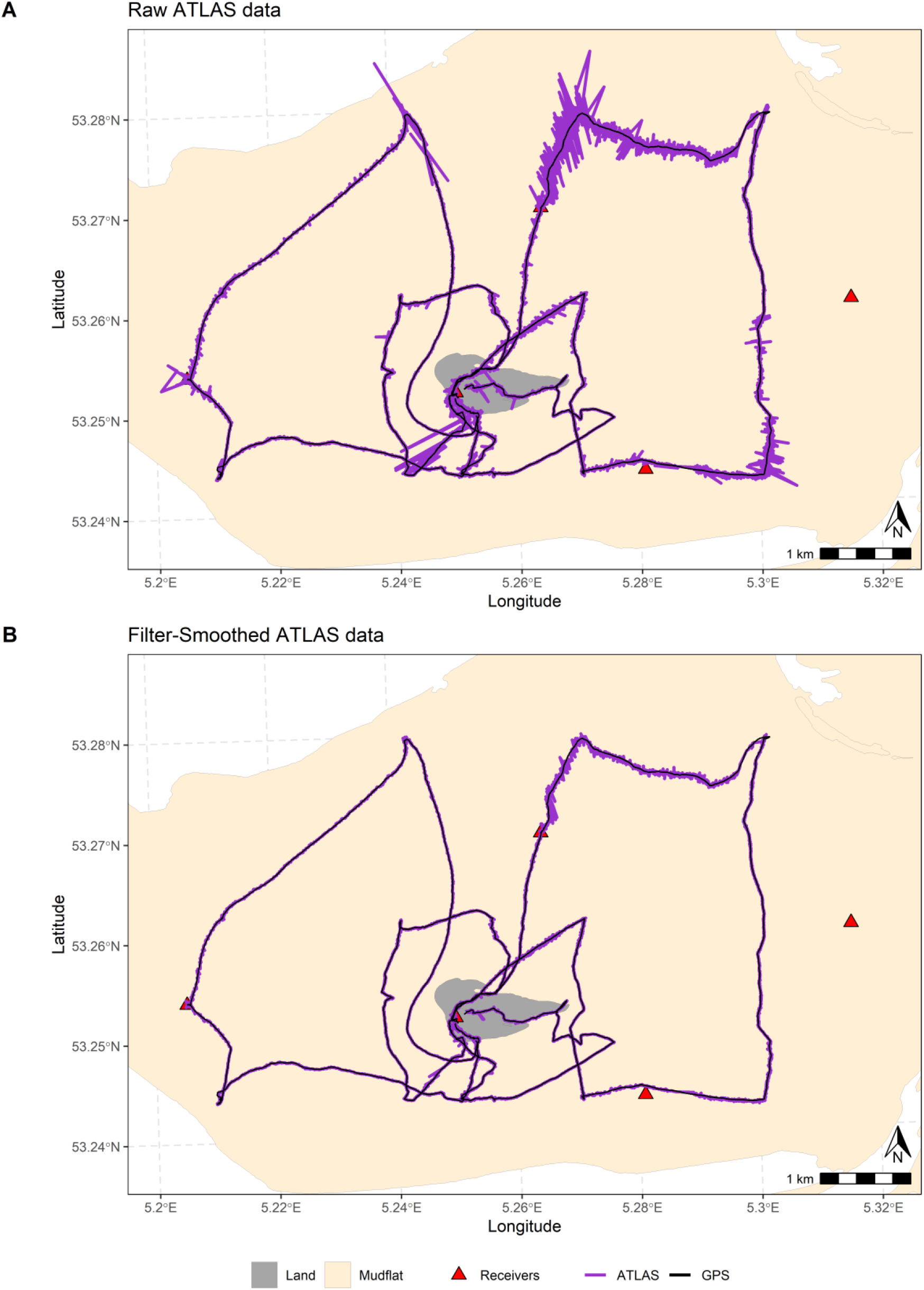
Comparison between ATLAS-(purple line) and GPS-(black line) derived localisations for raw (A) and filter-smoothed (B) data from the mobile test.

**Figure 5.**
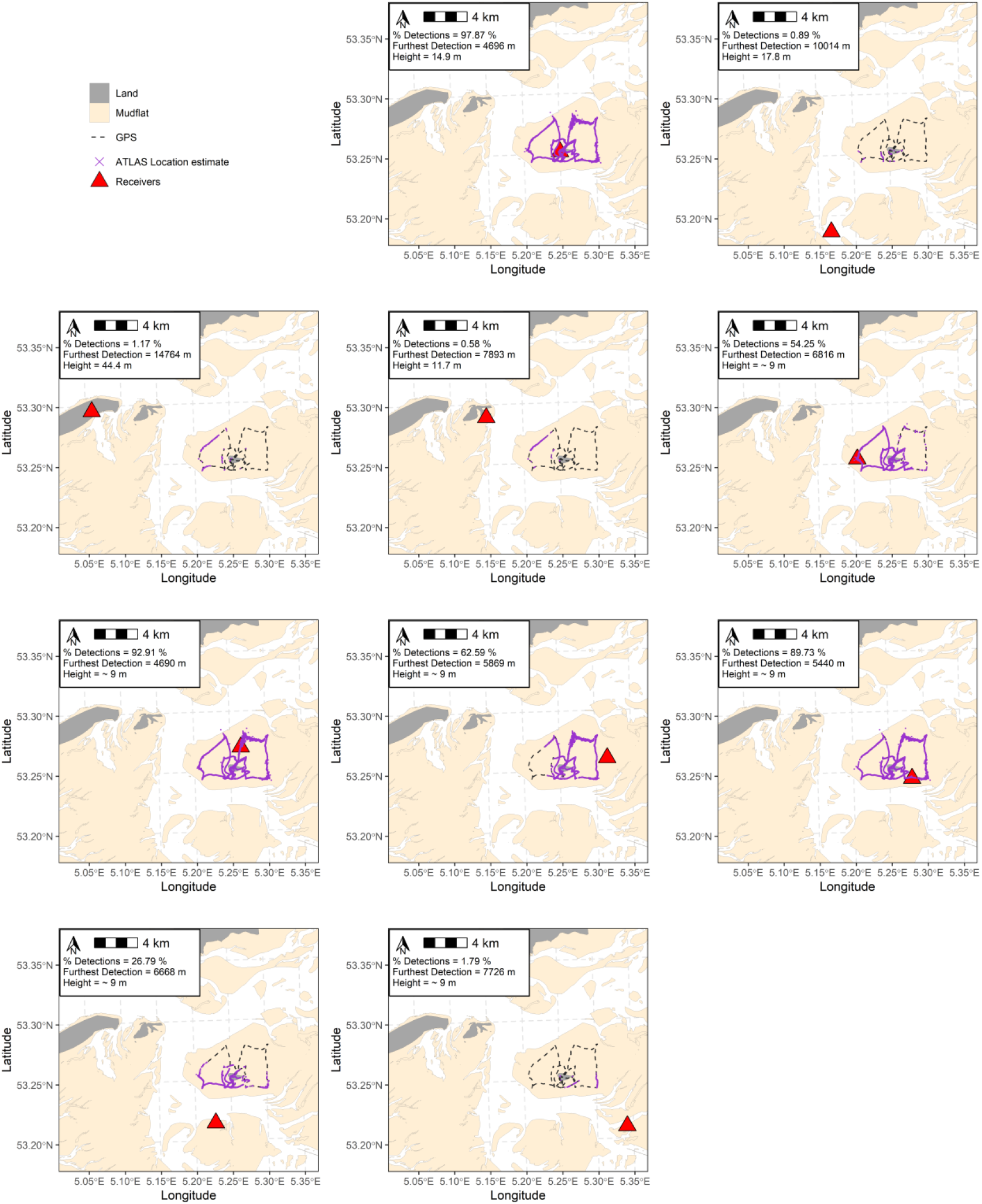
Detections for each of the contributing receiver stations and information on the percentage of total localisations that the receiver detected, the furthest detection from the receiver and the height of the receiver. Purple crosses denote location estimates that each receiver (red triangle) contributed to. The grey dashed line shows the path (tracked by GPS) of the researcher during the tests.

## Discussion

ATLAS location estimates were comparable to GPS. The median difference between raw ATLAS-derived location estimates and GPS-derived location estimates was 4.2 m for stationary tags and 5.7 m for moving tags. Accuracy was higher if more receivers detected the tag. However, the tag’s location in respect to the receiver array configuration also had a large effect, with less accurate estimates occurring when the tag was on the outskirts or outside the array of receivers. More accurate ATLAS localisations can therefore be achieved through strategic placing of receiver stations. We show that errors can be mitigated through a simple filter-smoothing process, as is routinely and intensively applied to raw GPS location estimates (Kaplan & Hegarty, 2005). Thus, ATLAS provides a viable and accurate alternative to GPS for regional-scale systems.

In TOA systems, error scales as a function of array geometry and location estimates are generally less reliable outside the array (MacCurdy et al., 2018). We found the most extreme errors occurred when the tag outside of the receiver array, matching the results seen in another ATLAS system (Beardsworth, 2020). Specifically, for the sections of the study that occurred in the most northern part of the mudflat, all detecting receiver stations were located to the south of the tag, outside the core array of receiving stations. While one receiver station was situated to the north of Griend, on Terschelling, 10 km away, this seems to be outside the range of reception, despite another receiver to the West, on Vlieland, detecting the tag 14.8 km away. The range at which receivers can detect transmissions can differ markedly between habitat types and topography, therefore this is highly site dependent (MacCurdy et al., 2009). For instance, detections of an ATLAS tag have been previously shown to differ over short distances due to the signals being blocked by hills (Beardsworth, 2020). However, the WATLAS system has very few topographical obstacles that limit reception of transmissions by receivers. It is therefore likely that other factors, such as height of the tag and/or receiver play an important role in reception.

Signal detection requires a clear ‘line of sight’ between the receiver and the tag. Because of the effects of reflections from ground and sea, higher receiver stations can typically continue detecting tags at larger distances than receivers that are closer to the earth’s surface (Xia et al., 1993). This may also explain why the highest receiver station (44.4 m high on Vlieland) was able to detect the tag almost 15 km away, but the Terschelling receiver station (35.1 m) was unable to detect the tag 10 km away. While receiver height is important, tag height also affects the reception of a signal. Tags attached to animals in flight can have much larger detection ranges than animals on the ground (Xia et al., 1993); for instance, localisations of Egyptian fruit bats (*Rousettus aegyptiacus*) during flight were based on receiver detections from up to 40 km away in the Hula ATLAS system (Toledo et al., 2020). If flight locations (as opposed to ground locations) are sufficient for a particular study, a sparser array of receivers may be appropriate to maximise coverage of an area. Ground locations can instead be inferred from take-off and landing locations. Similar inferences are made in marine systems, where diving behaviour prevents satellite tracking, thus movement tracks are reconstructed between surfacing events (Hussey et al., 2015). For systems that track ground-dwelling animals, receivers should be positioned <5 km away from the study area as we (with the tag 1.2 m above the ground) found the most successful receivers (>90% of signals detected) were within 5 km from the test locations.

While we did not specifically assess signal strength in this study, this can also have a strong influence on the accuracy of a location estimate. Signal strength can be affected by multiple causes. For instance, the height of the receiver’s antenna, the height of the tag antenna and the distance between them influences the strength of the signal at the receiver. Another factor that affects the signal strength is the transmitter power. While the transmitters in ATLAS tags have a power of 10mW, the relatively short tag antenna (which is necessary for use with animal-borne tags) reduces efficiency. Ping duration (8ms) and bandwidth (1MHz) also influence accuracy (Weller-Weiser et al., 2016). However, these values are typical for ATLAS and are optimized for miniature tags. Finally, interference from other transmitters, such as radio remote controls, can degrade the performance of the system. Due to the remoteness of the Wadden sea, there is little interference in the 434MHz band. This is evident from the fact that the gain, moderated by automatic gain control (AGC) at each receiver, is often very high, unlike in heavily populated areas. Nevertheless, this level of interference is site dependent and should be considered when establishing an ATLAS system.

The number of receivers detecting a tag also influenced accuracy. Tag signals detected by ≥ 4 receivers had a median error of ≤ 5.6 m while signals detected by 3 receiver stations had a median of 10.7 m error. This result may tempt users of ATLAS systems to use the number of receivers that contribute to a location estimate to filter movement data, removing 3-receiver localisations. Indeed, this has previously been suggested (Weller-Weiser et al., 2016). However, despite ten (out of a potential 26) receiver stations detecting the tag at various positions on the mudflat, 22.8% of signals were detected by only three receivers. Implementing such a broad filter may be detrimental to the amount of data retained about the animal’s movement. Furthermore, while an error of 10.7 m is acceptable in many cases, our simple filter-smoothing process reduced the error of 3-receiver location estimates to 6.3 m. We refer the reader to Gupte et al. (2020) for an in-depth assessment of filtering and smoothing techniques for ATLAS data.

It is important to note again that throughout this study, we based measures of accuracy on GPS location estimates yet GPS itself is susceptible to errors (Hofman et al., 2019). According to its specification, our particular GPS unit gives location estimates of <10 m error, therefore it is possible that ATLAS could be more or less accurate than this. For studies with no true location with which to measure accuracy, standard deviation in fixed locations gives an indication of the precision of a system. The standard deviation for 10 of the 16 stationary test sites was <5 m and thus, apparently similar to GPS (Forin-Wiart et al., 2015; Tomkiewicz et al., 2010). After filter-smoothing, 12 of the 16 sites had a standard deviation of <5 m. In these locations, the mean number of receivers detecting the tag signal ranged from 3.8-5.8 but all locations were central in the array, indicating that arrangement of receivers may be more important than concentration of receivers. Even the least accurate test location (site 2: which was outside the receiver array and detected by only a mean of 3.4 receiver stations), had a standard deviation of 15.1 m after filter-smoothing. Since most environmental covariates are measured at larger scales than this, ATLAS gives an appropriate level of accuracy and precision for many studies.

In summary, we provide evidence for the high accuracy and precision of a relatively novel localising system, ATLAS. In the design of an ATLAS system, we suggest that receiver stations are placed as high as possible and surround the study site with a direct line of sight. The density of receivers is important but depending on the number of receivers available for a study, trade-offs may have to be made between coverage and density of receivers. Considerations should also be made according to the height of the tag, with low-ground dwelling species requiring a denser receiver array than species that may fly between places of interest. ATLAS provides an opportunity to track animals remotely at high spatial and temporal resolution that rivals GPS technology at regional-scales and can be effective even with only 3 receiver stations. While we focus our study on a flat, intertidal region, ATLAS is not limited to flat landscapes. Several ATLAS systems have been operating successfully in hilly landscapes and complex agricultural systems (all other ATLAS systems), as well as wetlands (Israel) and woodlands (UK). These systems have successfully tracked >50 species of birds, mammals and reptiles, including small (8-15g) insectivorous bats and passerine birds for which GPS tracking at high temporal resolution is practically infeasible. In the focal ATLAS system, two smaller-bodied bird species, sanderling, *Calidris alba* (∼50 g) and red knot, *Calidris canutus* (∼120g) have been successfully tracked. Approximately ∼200 individuals per year are tagged, enabling the monitoring of complex biotic and abiotic interactions. However, it must be noted that ATLAS is limited to much smaller scales than GPS and therefore is unsuitable for certain studies. Furthermore, unlike military and commercially motivated multipurpose development of GPS, the ATLAS system was developed and is being applied by multidisciplinary teams of scientists for movement ecology research (Toledo et al., 2016). Each ATLAS system has been installed and is maintained by scientists, and often requires collaborations with landowners and organisations for establishing receiver locations. This is in stark contrast to (often) plug-and-go GPS tags. However, the advantages of ATLAS are large and we encourage further use of this high-throughput wildlife tracking system (contact S. Toledo and R. Nathan) which is suitable for use with a broad range of study systems that require accurate, high-throughput tracking to answer biological questions.

## Acknowledgements

This work was partly funded by the Dutch Research Council grand VI.Veni.192.051 awarded to AIB. We are grateful to the Minerva Center for Movement Ecology for funding ATLAS development and implementation and to the core Israeli ATLAS team (Yotam Orchan, Yoav Bartan, Oren Kishon, Adi Weller-Weiser, Ingo Schiffner, Anat Levi and Sivan Margalit) and its main practitioners (Motti Charter, David Shohami, Yosef Kiat, Emmanuel Lourie, Nadav Ganot, Aya Goldshtein and Rea Shaish) for their valuable help in developing ATLAS over the past 8 years. Many people and organisations are involved in hosting the ATLAS equipment to complete this study, without whom it would be impossible. We would therefore like to thank the following (in no particular order): Hoogheemraadschap Hollands Noorderkwartier, Koninklijke Nederlandse Redding Maatschappij, Staatsbosbeheer, Marine Eco Analytics, Koninklijke Luchtmacht, Het Posthuys, Natuurmonumenten, Wetterskip Fryslan, Afsluitdijk Wadden Center, Vermilion, Rijkswaterstaat, Carl Zuhorn, Lenze Hofstee and Lydia de Loos. We would also like to thank the RV Navicula and RV Stern staff and volunteers for facilitating the work around Griend and Anita Koolhaas, Hinke and Cornelis Dekinga and Job ten Horn for their help with building the receiver stations. We thank Selin Ersoy, who proofread the manuscript and gave constructive feedback. Finally, we thank Joah Madden and Luca Borger, whose constructive conversations and comments on CEB’s PhD thesis inspired the writing of this manuscript.

## Authors’ contributions

CEB and AIB conceived the idea for the manuscript; CEB, EG and AIB collected the data; CEB and EG analysed the data; CEB led the writing of the manuscript; RN and ST developed the ATLAS system and provided support throughout data collection; AIB, BD, AD and FvM built and maintained the WATLAS system and tags, and provided local support throughout data collection; all authors contributed critically to the drafts and gave final approval for publication.

## Data accessibility

Data and relevant code for this research are stored in GitHub: https://github.com/CBeardsworth/watlas_validation and have been archived within the Zenodo repository: https://doi.org/10.5281/zenodo.4527613

