## Appendix S1 for "Validating a high-throughput tracking system: ATLAS as a regional-scale alternative to GPS"

### Electronic Supplementary Material for: Validating a high-throughput tracking system: ATLAS as a regional-scale alternative to GPS

Christine E. Beardsworth*, Evy Gobbens, Frank van Maarseveen, Bas Denissen, Anne Dekinga, Ran Nathan, Sivan Toledo, and Allert I. Bijleveld

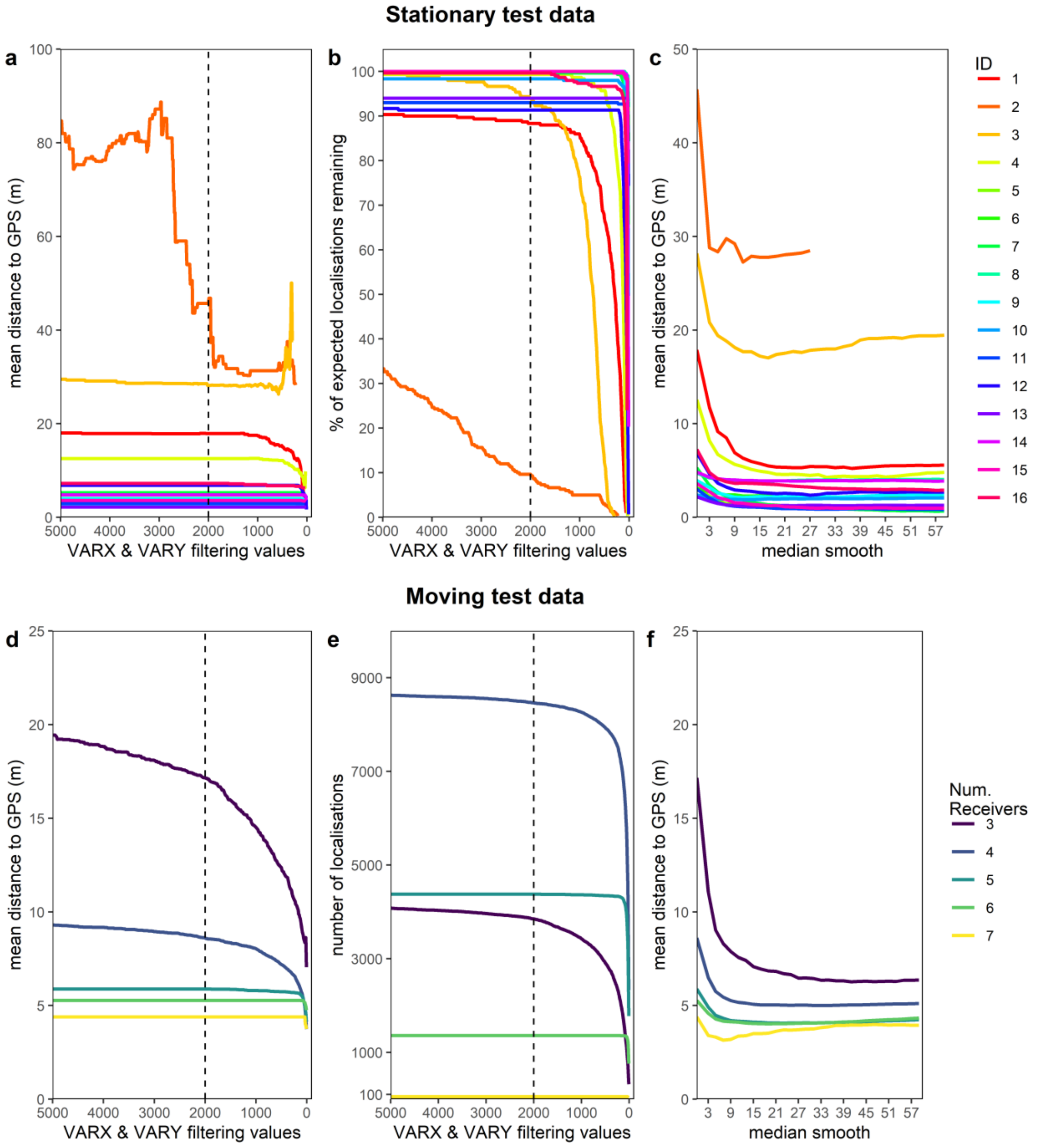


**Appendix S1. Filtering and smoothing efficacy.** The use of ATLAS-derived accuracy estimates increased the accuracy of the most erroneous stationary site (a). We used a threshold of 2000 VARX and VARY (dashed vertical line) in this study and found that this filter only had an effect on the least accurate site. Setting this threshold begins a trade-off between accuracy (a) and retaining localisations

(b) which can then further be improved through smoothing (c). Using the moving test data, we also found a similar trade-off between accuracy (d) and the number of localisations retained (e), although VARX and VARY had little effect on the accuracy of 5,6 and 7 receiver-derived localisations (d). Again, these location estimates were further improved by using a median smoother (f).
